# Depth-dependent PSF calibration and aberration correction for 3D single-molecule localization

**DOI:** 10.1101/555730

**Authors:** Yiming Li, Yu-Le Wu, Philipp Hoess, Markus Mund, Jonas Ries

**Affiliations:** European Molecular Biology Laboratory (EMBL), Cell Biology and Biophysics unit, Meyerhofstr. 1, 69117 Heidelberg, Germany; Collaboration for joint PhD degree between EMBL and Heidelberg University, Faculty of Biosciences

## Abstract

3D Single molecule localization microscopy relies on fitting of the individual molecules with a point spread function (PSF) model. The reconstructed images often show local squeezing or expansion in z. A common cause are depth-induced aberrations in conjunction with an imperfect PSF model calibrated from beads on a coverslip, resulting in a mismatch between measured PSF and real PSF. Here, we developed a strategy for accurate *z*-localization in which we use the imperfect PSF model for fitting, determine the fitting errors and correct for them in a post-processing step. We present an open-source software tool and a simple experimental calibration procedure that allow retrieving accurate *z*-positions in any PSF engineering approach or fitting modality, even at large imaging depths.

## 1. Introduction

3D single molecule localization microscopy (SMLM) is a powerful technology for cell and molecular biology as it can resolve biological structures with nanoscale resolution in all three dimensions [1]. In most approaches, the *z*-positions of single fluorophores are determined from their shape, either by introducing additional aberrations [2,3] or by evaluating two or more channels focused at different focal planes [4] to increase their *z*-dependence. To precisely relate shape parameters to absolute *z*-positions or to determine an experimental 3D point-spread function (PSF) model [5,6], an accurate calibration is required. Commonly, this consists of acquiring a *z*-stack of beads immobilized on a coverslip. An intrinsic assumption is that the PSF on the coverslip approximates the one deep in the sample. However, this is not generally valid due to depth-induced aberrations, especially when the refractive indices of immersion oil and sample medium are different. Most researchers use oil objectives for SMLM, because of their high collection efficiency, compatibility with total internal reflection excitation and simple implementation of focus stabilization. However, the refractive index mismatch between the glass and the sample leads to strong, mostly spherical aberrations when imaging above the coverslip inside the sample. These aberrations are depth-dependent and can be substantial even for moderate imaging depths of a few micrometers, resulting in systematic errors in the *z*-localization. Furthermore, when imaging beads on coverslips in aqueous solutions while using a high numerical aperture oil objective (e.g. NA = 1.45, *n*_oil_ = 1.52), supercritical angle fluorescence occurs that further deviates the PSF on the coverslip from that deep inside the medium [7].

A mismatch between the calibrated PSF model and the real PSF within the biological specimen often leads to a distortion of the reconstructed super-resolution images. To compensate for these aberration-induced localization errors, several techniques have been proposed, either by active aberration correction with adaptive optics [8,9] or realistic PSF estimation by numerical computation and experimental PSF measurements. Since dedicated optics and expertise are needed for implementation of adaptive optics, active aberration correction is still only used in few expert labs. For numerical computation, both theoretical PSFs [10] and a phase retrieved PSF [11–13] have been used to model the PSF at different depths. However, these methods suffer from imperfect optics and complicated data analysis. Alternatively, the 3D depth-dependent PSF can be directly determined experimentally, e.g. using an optical trap, which is cumbersome and requires dedicated equipment [14]. A common strategy to generate a depth-dependent PSF calibration is to measure many fluorophores at different distances away from the coverslip to generate a calibration curve. This can be done either by 1) embedding beads in a gel at different heights [15] or 2) attaching fluorophores on a defined curved surface [16]. Then, each fluorescent molecule gives a random sampling of the *z*-position. This approach works well for astigmatism-based 3D methods, because the calibration curve can be easily determined based on the PSF width in the x and y direction, respectively. For more complex PSFs [1], such kind of calibration curve might not be easily generated. Here, we develop a new approach and a simple open source software tool to correct for depth-induced fitting errors for any PSF-engineering approach and any fitting modality. Our approach is to take many *z*-stacks of beads immobilized in a gel above the coverslip and fit them with a PSF model of choice to obtain the fitted *z*-position in dependence on the objective position. The correction for *z*-positions can be determined at any depth by interpolating between different beads and is applied to an SMLM data set in a post-processing step.

## 2. Depth-dependent *z*-calibration

A schematic of the procedure for calibrating the depth-dependent *z*-correction is shown in Fig. 1. To calibrate the magnitude of *z*-localization errors for a specific PSF model in dependence on the imaging depth, we embedded fluorescent beads in a layer of agarose gel above the coverslip and acquired *z*-stacks at many objective positions in a range from 1 µm below to several micrometers above the coverslip with a spacing of typically 20 nm to 50 nm (Fig. 1a). Each frame in the stack corresponds to a different focal plane. As an estimate for the *z*-position of a bead relative to each focal plane, we first determined the frame in the stack for which the fitter returned a *z*-position of zero as the focal position (z = 0). By interpolation, we could achieve this with an accuracy better than the distance of the calibration planes. This measure for the *z*-position of a bead is well-suited for astigmatism-based 3D SMLM: as depth-induced aberrations are mostly symmetric, they do not change the asymmetry of the PSF dramatically. Thus, the focal plane, in which a bead appears symmetric is largely independent of the aberrations and thus can be defined as its nominal bead position, which can be directly read from the objective positions.

**Fig. 1.**
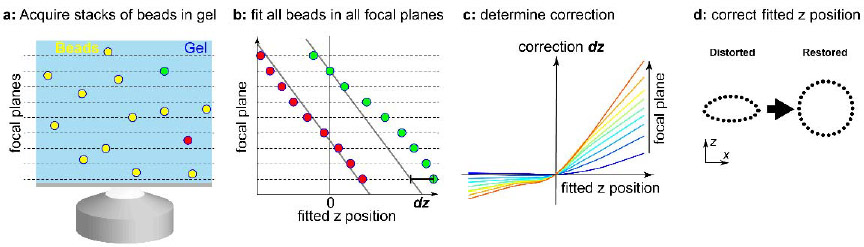
Schematic of the approach to calibrate the depth-dependent *z*-correction. (a) Fluorescent beads are embedded in agarose gel at random depths and are imaged at different focal planes. (b) Example of fitted *z*-position versus nominal focal plane position (objective position from piezo stage) for two beads at different heights (red and green in a). Due to aberrations, the fitted position is not linear with the nominal z position (focal plane). This allows us to determine the correction *dz* in dependence on the focal plane. (c) Calibration curve showing the correction *dz* of fitted *z*-values in dependence on the fitted *z*-values and the position of the focal plane above the coverslip. (d) In an SMLM experiment, the data is fitted with the same PSF model as the beads in gel, and the fitted *z*-positions of the fluorophores are corrected using the calibration from (c).

We can then determine the nominal *z*-position of each bead at each frame and compare it with the fitted *z*-position (Fig. 1b). As we take into account many bead stacks, we can determine this *z*-correction for many combinations of fitted *z*-positions and focal plane positions (Fig. 1c). We then performed a robust interpolation of these data with a smoothed cubic B-spline by iteratively removing outliers with a too large distance from the smoothed surface, until no outliers were present any longer. This calibration is then used as a look up table to correct measured *z*-positions in an SMLM experiment. For this, one needs to determine the approximate focal position above the coverslip. This can be achieved for instance by focusing first on dyes unspecifically bound to the glass or residual fluorescence on the glass surface, before focusing to the area of interest and reading out the difference in objective depth. Once we know the objective depth, the interpolated *z*-correction function directly returns the *z*-correction from the focal plane position and the fitted *z*-values (Fig. 1d). To account for the difference between the objective position and actual *z*-position due to refractive index mismatch, we further multiply the corrected *z*-values by a refractive index mismatch factor of in our case 0.8 [2].

## 3. Material and Methods

### 3.1 Optical Setup

Experiments were performed on a customized microscope [17] equipped with a high NA oil immersion objective (160x, 1.43-NA oil immersion, Leica, Wetzlar, Germany) at room temperature (24 °C). A laser combiner (LightHub®, Omicron-Laserage Laserprodukte, Dudenhofen, Germany) with Luxx 405, 488 and 638, Cobolt 561 lasers was used for SMLM image acquisition. The lasers were triggered using a FPGA (Mojo, Embedded Micro, Denver, CO, USA) allowing microsecond pulsing control of lasers. The lasers are first passed through a speckle reducer (LSR-3005-17S-VIS, Optotune, Dietikon, Switzerland), and then guided through a multimode fiber (M105L02S-A, Thorlabs, Newton, NJ, USA). The output of the fiber is first magnified by an achromatic lens and then imaged into the sample. A laser clean-up filter (390/482/563/640 HC Quad, AHF, Tübingen, Germany) is placed in the beam path to remove any fluorescence generated in the excitation path. A focus lock system was implemented using the signal of a near-infrared laser reflected by the coverslip and its detection by a quadrant photodiode which was controlled in close-loop. The focus can be stabilized within ±10 nm over several hours [18]. In the emission path, the fluorescence was filtered by a bandpass filter (700/100, AHF) and recorded by an EMCCD camera (Evolve512D, Photometrics, Tucson, AZ, USA). Typically, we acquire ~200,000 frames with 15 ms exposure time until all the fluorescent molecules are bleached. The excitation laser power was ~15 kW/cm^2^. The pulse length of the 405 nm laser is automatically adjusted to retain a constant number of localizations per frame.

### 3.2 Data Processing

All single molecule image data was analyzed with an experimental PSF model calibrated by 100 nm TetraSpeck fluorescent beads (T7279, ThermoFisher Scientific) immobilized on the coverslip as described before [5]. Briefly, the bead images were firstly averaged across different fields of view. The averaged 3D bead stack was then interpolated by cubic spline interpolation. Here, the 64 spline coefficients were calculated for each voxel. Finally, the cubic spline PSF model was built based on these spline coefficients.

All the returned *x*, *y*, *z*-positions were corrected for residual drift by a custom algorithm based on redundant cross-correlation [19]. Briefly, the localizations were binned in typically ten time windows and individual super-resolution images at each time window were reconstructed. The pair-wise image cross-correlation of all images with all others was determined and fitted with a spline interpolation. The lateral drift trajectory was calculated and corrected accordingly. For the 3D drift correction, we first performed a 2D drift correction and then applied a *z*-drift correction based on redundant 1D cross-correlations.

For calibrating the 3D aberrations at different depths, the 3D bead stacks in gel are fitted to the imperfect PSF model on coverslip. The position of the coverslip was determined by the beads with lowest nominal focal position. The nominal position of each bead was determined by the objective position when the fitted *z*-position is 0. For each bead, the fitted *z*-positions as a function of each nominal focal plane were plotted. The correction value was then calculated based on the difference between the fitted *z*-positions and its nominal *z*-positions at each focal plane. A 2D smoothed spline surface was generated on these correction values. Any points that are too far away from the surface are eliminated. The software for generating the depth dependent 3D aberration curve is open source. The instructions and an example of its use are available on Github https://github.com/jries/fit3Dcspline.

To quantitatively analyze the distance between the two rings of the nuclear pore complex protein Nup107, we fit a model consisting of two parallel rings to the localizations of each nuclear pore complex using maximum likelihood estimation. By fitting the data, we corrected the tip and tilt of each individual nuclear pore complex to improve the precision when calculating the distance of the 2 rings from the z intensity profile.

### 3.3 Embedding of fluorescent beads in agarose gels for calibration measurements

We prepared a 1% [w/v] solution of low melting point agarose (A9414, Sigma-Aldrich) in water, heated it up to completely dissolve the agarose, and let it cool down to approx. 40 °C. We then vigorously vortexed the stock solution of TetraSpeck fluorescent beads, and added 2.5 µL to 400 µL of the agarose solution, and vortexed again. We then added a 50 µL drop onto a coverslip. After a few minutes, we mounted the sample in water, and imaged the ~1 mm thick gel that contained immobilized fluorescent beads throughout.

### 3.4 Biological Sample Preparation

Round 24 mm high precision glass coverslips No. 1.5H (117640, Marienfeld, Lauda-Königshofen, Germany) were used for all experiments. Coverslips were cleaned overnight in a 1:1 mixture of concentrated HCl and methanol, rinsed with millipore water until neutral, dried and UV sterilized in a standard cell culture hood before cell culture.

For imaging of clathrin-coated pits, SK-MEL-2 cells (kind gift from David Drubin, described in Ref. [20]) were cultured under adherent conditions in DMEM/F-12 (Dulbecco’s Modified Eagle Medium/Nutrient Mixture F-12) with GlutaMAX, phenol red (ThermoFisher 10565018) supplemented with 10% [v/v] FBS (ZellShield™, Biochrom AG, Berlin, Germany), and 30 mM HEPES at 37°C, 5% CO_2_ and 100% humidity. Cells were fixed using 3% [w/v] formaldehyde (FA) in cytoskeleton buffer (CB; 10 mM MES pH 6.1, 150 mM NaCl, 5 mM EGTA, 5 mM D-glucose, 5 mM MgCl_2_, described in Ref. [21]) for 20 minutes. Fixation was stopped by incubation in 0.1% [w/v] NaBH_4_ for 7 minutes. The sample was washed with PBS three times, and subsequently permeabilized using 0.01% [w/v] digitonin (Sigma-Aldrich, St. Louis, MO, USA) in PBS for 15 minutes. After washing twice with PBS, the sample was blocked with 2% [w/v] BSA in PBS for 60 minutes, washed again with PBS, and stained for 3-12 hours with anti-clathrin light chain (sc-28276, Santa Cruz Biotechnology, Dallas, TX, USA, diluted 1:300) and anti-clathrin heavy chain rabbit polyclonal antibodies (ab21679, Abcam, Cambridge, UK, diluted 1:500) in 1% [w/v] BSA in PBS. The sample was washed with PBS three times, and incubated with a donkey anti-rabbit secondary antibody (711-005-152, Jackson ImmunoResarch, West Grove, PA, USA), which was previously conjugated with Alexa Fluor 647-NHS at an average degree of labeling of 1.5, for 4 hours. Finally, the sample was washed three times with PBS prior to imaging.

For imaging of nuclear pore complexes, genome-edited U-2 OS cells that express Nup107-SNAP [5,22] were cultured under adherent conditions in Dulbecco’s Modified Eagle Medium (DMEM, high glucose, w/o phenol red) supplemented with 10% [v/v] FBS, 2 mM L-glutamine, non-essential amino acids, ZellShield™ (Biochrom AG, Berlin, Germany) at 37 °C, 5% CO_2_ and 100% humidity. All incubations were carried out at room temperature. For nuclear pore staining, the coverslips were rinsed twice with PBS and prefixed with 2.4% [w/v] FA in PBS for 30 seconds. Cells were permeabilized with 0.4% [v/v] Triton X-100 in PBS for 3 minutes and afterwards fixed with 2.4% [w/v] FA in PBS for 30 minutes. Subsequently, the fixation reaction was quenched by incubation in 100 mM NH_4_Cl in PBS for 5 minutes. After washing twice with PBS, the samples were blocked with Image-iT™ FX Signal Enhancer (ThermoFisher Scientific, Waltham, MA, USA) for 30 minutes. The coverslips were incubated in staining solution (1 µM benzylguanine Alexa Fluor 647 (S9136S, NEB, Ipswich, MA, USA); 1 mM DTT; 1% [w/v] BSA; in PBS) for 50 minutes in the dark. After rinsing three times with PBS and washing three times with PBS for 5 minutes, the sample was mounted for imaging.

For dSTORM imaging, coverslips were mounted in 500 µL blinking buffer (50 mM Tris pH 8, 10 mM NaCl, 10% [w/v] D-glucose, 35 mM 2-mercaptoethylamine (MEA), 500 µg/mL GLOX, 40 µg/mL catalase, 2 mM cyclooctatetraene (only for imaging of clathrin-coated pits)).

## 4. Experimental Results

Firstly, we validated our approach on beads (Fig. 2). We embedded 100 nm fluorescent beads in an agarose gel and acquired typically tens of *z*-stacks at different fields of view from 1 µm below to 5 µm above the coverslip with a step size of 20 nm. Each bead represents a random sampling of the *z*-position in each focal plane. In each frame, all the beads were fitted with a PSF model that was calibrated on the coverslip. The fitted *z*-positions were plotted against the objective positions (Fig. 2a). The distance of the beads from the coverslip can be directly read from the objective position when the fitted z is 0. The position of the coverslip is determined similarly with the beads attached to the surface of the coverslip. If the real PSF model is close to the PSF model calibrated on the coverslip, the fitted *z*-position would correspond to the objective position. This was true for beads less than ~500 nm away from the coverslip (Fig. 2a). However, for beads further away from the coverslip, a substantial correction of the fitted *z*-position is required to agree with the real *z*-position (Fig. 2b). Due to aberrations, a linear rescaling factor [10] is not enough for accurately correcting the fitted z as shown in Fig. 2b. Therefore, we interpolated these correction values across different focal plane and z positions with a smoothed cubic B-spline. To validate the effectiveness of these correction values, we applied them to all the beads in the 3D stacks. Before correction (left panel, Fig. 2c), the fitted *z*-positions are not equal to the distance of the bead from the focal plane (objective position in the current frame minus objective position when the fitted z is 0). The root mean square (rms) error is 148 nm and Pearson correlation coefficient c=0.9848. After correction (Right panel, Fig. 2c) the fitted *z*-positions show a very high correlation with their distance from the focal plane. The rms error is only 18 nm with a Pearson correlation coefficient c=0.9993.

**Fig. 2.**
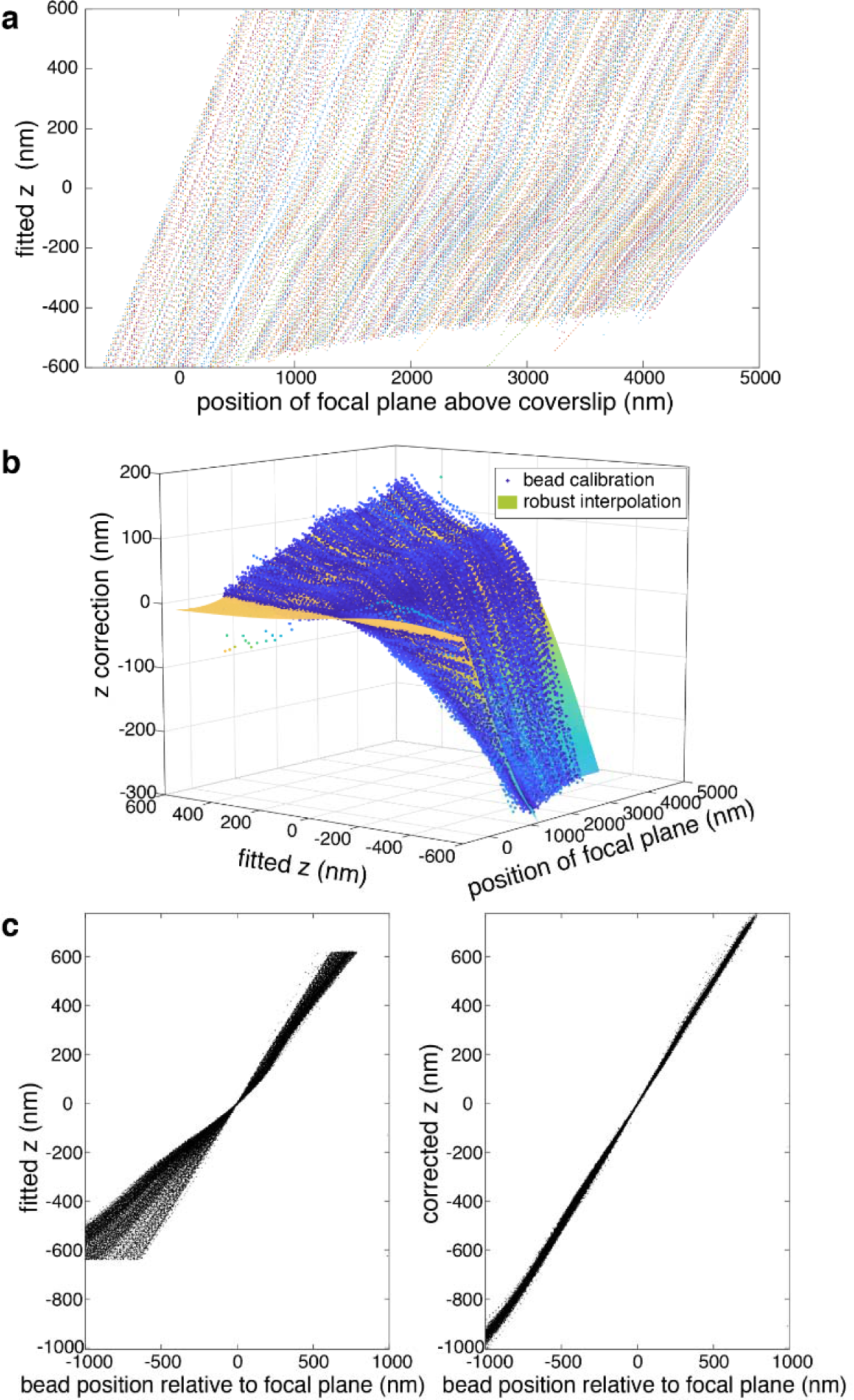
Correction of depth-induced aberrations. (a) 717 Beads, embedded in an agarose gel, were fitted with a PSF model that was calibrated on the coverslip. For deeper beads, the fitted *z*-position increasingly deviates from their distance from the focal plane. (b) Correction for fitted *z*-values in dependence on the fitted *z*-values and the position of the focal plane above the coverslip. (c) The left panel shows the fitted *z*-positions in dependence on the distance of the bead from the focal plane, and for many beads these are not equal (root mean square (rms) error 148 nm, Pearson correlation coefficient c=0.9848). The right panel shows the corrected *z*-positions, which now show a very high correlation with their distance from the focal plane (rms error 18 nm, Pearson correlation coefficient c=0.9993).

We then demonstrated the performance of our approach on SMLM imaging of a biological structure, namely clathrin coated pits (CCPs) in fixed SK-MEL-2 cells. Clathrin is a molecular scaffold that plays a major role in the formation of the coated vesicles during endocytosis. The single molecule data were fitted with an experimental astigmatic PSF model determined from beads on the coverslip as described before [5]. For CCPs close to the coverslip, a simple scaling factor correction for refractive index mismatch could already recover nicely the spherical geometry of the CCPs (Fig. 3a). However, for CCPs on the upper cell membrane, which is about 2 µm above the coverslip, the reconstructed images show squeezing and distortions in z (Fig. 3b). This is due to a mismatch of the real PSF model inside the sample and the PSF model calibrated on coverslip. Therefore, we applied the correction value calibrated with the beads in gel as described before (Fig. 2). After correction, we could fully recover the spherical geometry of clathrin-coated pits (Fig. 3b), thus we corrected for aberration induced fitting errors at a depth of 2 µm.

**Fig. 3.**
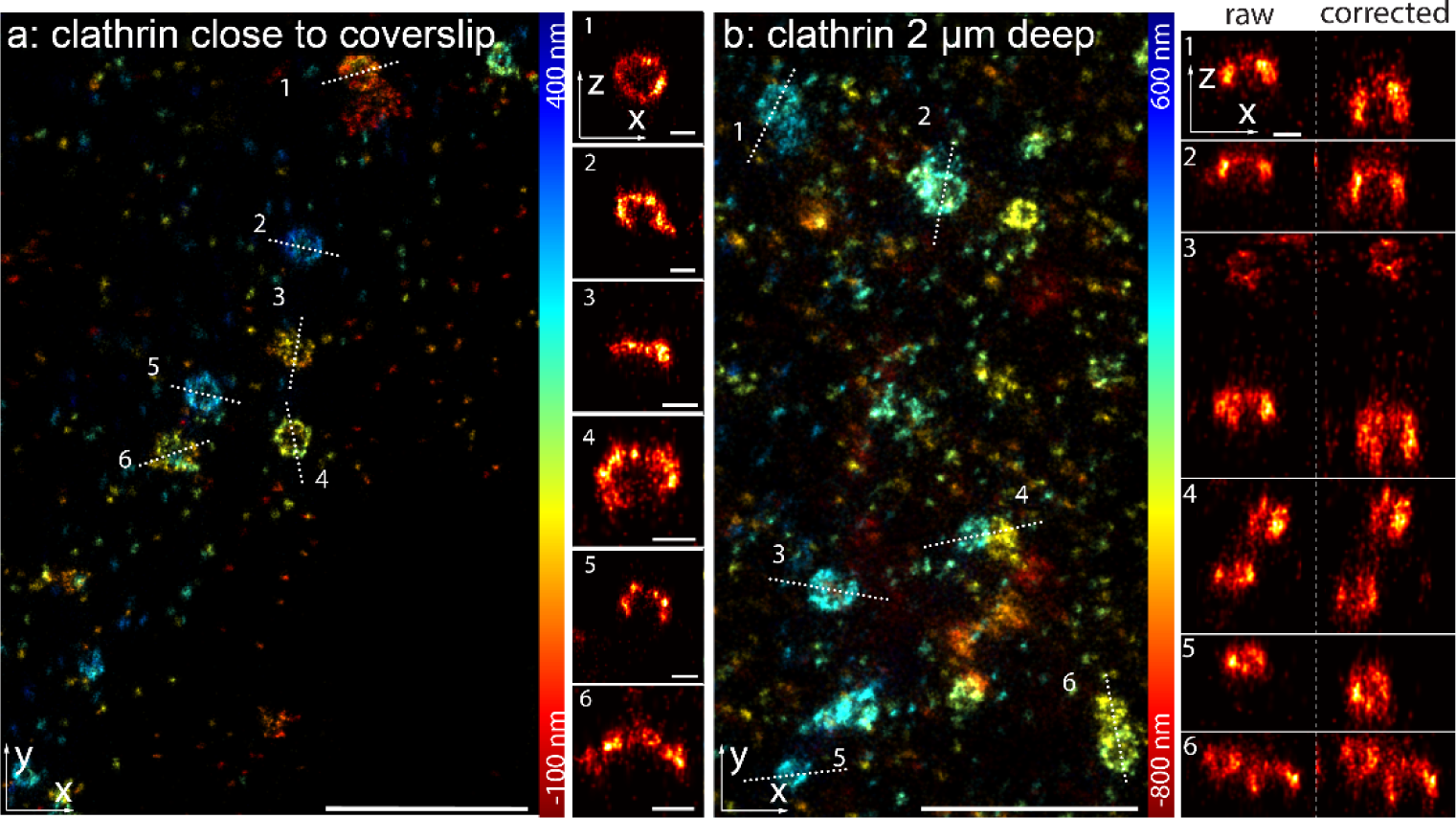
Imaging of clathrin-coated pits. (a) Clathrin-coated pits, close to the coverslip, immunolabeled with Alexa Fluor 647 conjugated antibodies, measured using dSTORM [23] and fitted with an experimental PSF determined from beads on the coverslip. (b) Clathrin-coated pits on the upper cell membrane, imaged 2 µm above the coverslip using an oil objective, show deformations in the side-view reconstructions. After correction of aberration-induced artifacts the spherical shape of the pits is recovered. Width of the line profiles: 50 nm. Scale bars: 1 µm for *x*-*y* reconstruction panels and 100 nm for *x*-z reconstruction panels.

We further tested our approach on a more challenging specimen by imaging the nuclear pore complex (NPC) protein Nup107 on the upper nuclear envelope which is about 5 µm away from the coverslip. Nup107 is part of the Y-shaped complex which forms a nucleoplasmic ring and a cytoplasmic ring, which can be resolved as two rings in SMLM when imaged close to the coverslip [5]. However, when we focused deep inside the cell and imaged on the top nuclear envelope, we found that the reconstructed image was squeezed (Fig. 4a). Due to the model mismatch of the PSF deep inside the cell and the one on the coverslip, most of the fitted positions were kept around the focal plane and showed a flattened surface (Fig. 4c). After correction (Fig. 4b and d) with the calibration curve of the beads in gel, we found that the fitted positions were not constrained around the focal plane anymore and a spherical surface of the nucleus envelop could be reconstructed.

**Fig. 4.**
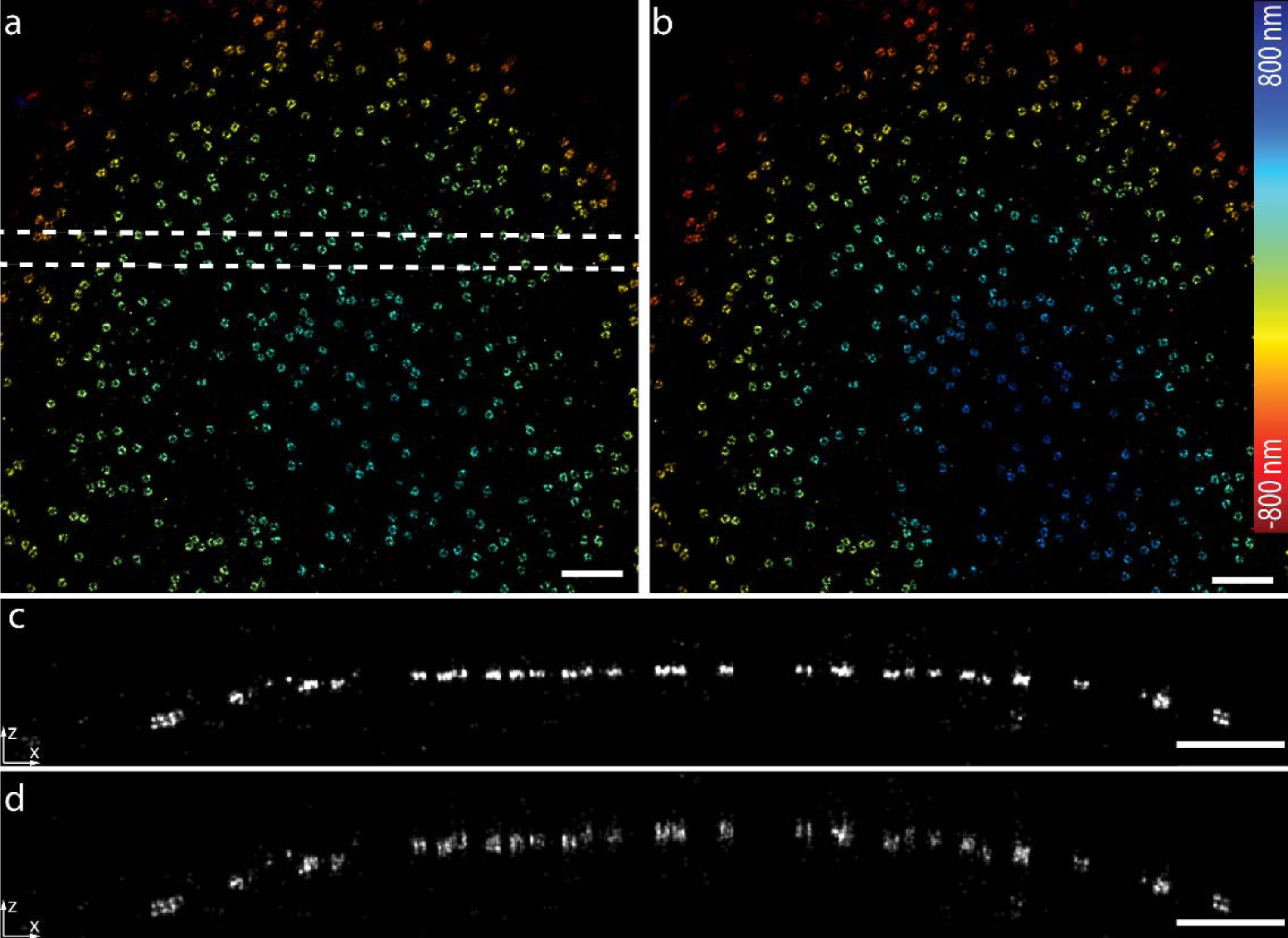
Imaging of nuclear pore complex protein Nup107 about 5 µm deep in the cell. (a) Nup107-SNAP-Alexa Fluor 647 imaged with dSTORM reconstructed by the PSF model obtained on the coverslip. (b) dSTORM image after applying the depth dependent z correction to a. (c) Side-view reconstruction of the region bounded by dashed line in a. (d) Side-view reconstruction of the same region as in c after applying the depth-dependent *z*-correction. Scale bars, 1 µm.

Finally, we applied our calibrated correction curve to the data close to the coverslip. Again, Nup107 was used to evaluate the effect of the correction (Fig. 5a, 340 nm above the coverslip). As expected, we could nicely resolve the two rings because the PSF at this depth is close to the one on the coverslip (Fig. 5 b and c). We determined the separation of the rings to be 56.5 (mean) ± 6.6 (SD) nm. To test for residual aberration-induced *z*-position errors, we plotted the distance between the two rings as a function of the central *z*-position of these two rings. Surprisingly, we found a strong z dependence with a Pearson correlation coefficient of 0.55 (Fig. 5d). This indicates that the PSF model mismatch could already play a significant effect even in such close proximity to the coverslip. Therefore, we corrected our data with the calibration curve as described before. After correction, the ring distances showed hardly any z dependence with a Pearson correlation coefficient of −0.03 (Fig. 5e). These results demonstrate that our method can effectively reduce the bias induced by the model mismatch between the real PSF and the one on coverslip even at depths close to coverslip.

**Fig. 5.**
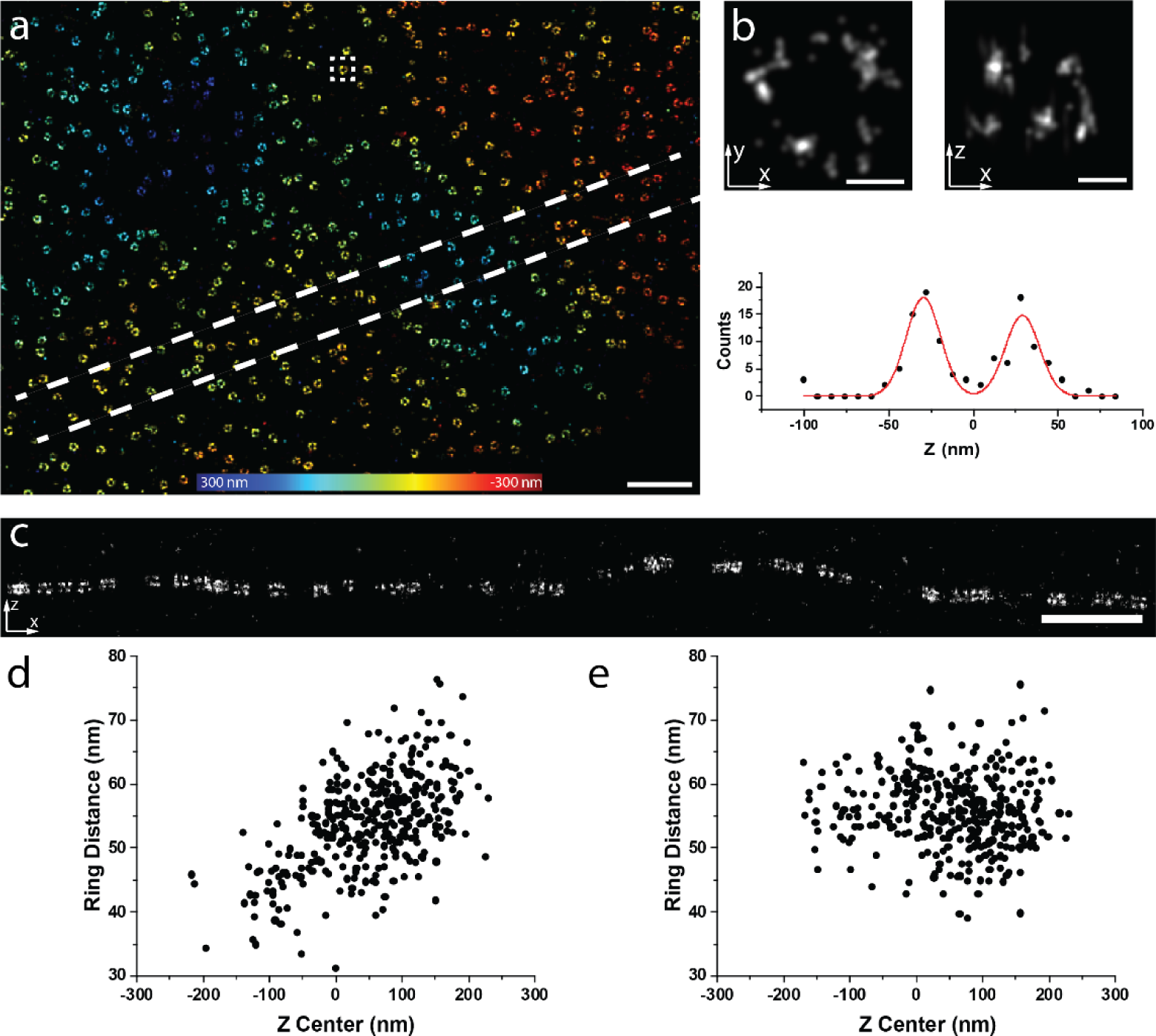
Imaging of Nup107 at 340 nm above the coverslip. (a) Nup107-SNAP-Alexa Fluor 647 imaged with dSTORM reconstructed by the PSF model obtained on the coverslip. (b) dSTORM image of a representative single NPC as indicated in the squared region in (a). Top images are *x*-*y* and *x*-z views. Bottom image is the intensity plot along z and a two Gaussian model was fitted on the data. (c) Side-view reconstruction of the region bounded by dashed lines in (a). (d) The distance of the two rings as a function of the central z position of each individual NPC before correction (Pearson correlation coefficient c=0.55). (e) The distance of the two rings as a function of the central z position of each individual NPC after correction (Pearson correlation coefficient c=-0.03). Scale bars, (a) and (c) 1 µm, (b) 50 nm.

## 5. Conclusions

The ability to accurately and easily calibrate the depth-dependent PSF is an important aspect to reconstruct accurate 3D super-resolution images deep in the specimen. However, it is complicated to directly measure the PSF deep in the sample. To overcome this problem, we developed an approach in which we fit the data using an imperfect PSF model calibrated from the beads on the coverslip and correct for the errors between the real and fitted *z*-positions in a post-processing step. This correction for depth-dependent axial distortions complements previous work correcting for depth-dependent lateral distortions [24]. For quantitative analysis, this correction was necessary already in the close vicinity of the coverslip, possibly because of how supercritical-angle fluorescence affects the bead-calibrated PSF model. Note that although our correction leads to a high accuracy, it does not improve the precision of *z*-position estimates as the localizations are still fitted with a sub-optimal PSF model.

Our approach to correct for depth-induced localization errors requires only a simple calibration step with beads embedded in gel, and as such uses only standard reagents readily available in most labs. It is compatible with any fitting software and does not require additional hardware. Together with our open-source software, this approach can be easily and readily applied in any lab, and thus broadly enables accurate 3D SMLM at depths previously only accessible to specialized labs.

